# Delivery of defective interfering RNA antivirals using hyperbranched poly(beta-amino ester) nanoparticles

**DOI:** 10.64898/2026.05.29.721911

**Authors:** Shun Yao, Jason Atkins, Priya Dhole, Ines Pena-Novas, Julien H. Arrizabalaga, Arun K. Sharma, Krishne Gowda, Daniel Hayes, Gabriella Worwa, Jens Kuhn, Marco Archetti

## Abstract

Hyperbranched poly(beta-amino ester) (hPBAE) nanoparticles represent a promising platform for nucleic acid delivery, particularly to the lungs. In this study, we evaluate the potential of hPBAE nanoparticles to deliver defective interfering RNA (diRNA) antivirals targeting betacoronaviruses under a range of formulations and storage conditions. hPBAE–diRNA nanoparticles demonstrated efficient cellular uptake of functional diRNA across diverse cell types, conferred protection against nuclease-mediated degradation, and exhibited low in vitro cytotoxicity. In vivo, these nanoparticles enabled effective delivery of functional diRNA to the lungs of golden hamsters without inducing adverse physiological effects. Collectively, these findings support hPBAE nanoparticles as a safe and effective platform for diRNA delivery for the treatment of respiratory viral infections.

Graphical Abstract.
Defective interfering RNA was mixed with hyperbranched poly(beta-amino ester) nanoparticles and delivered to cells in vitro and to golden hamsters in vivo, to measure toxicity and the replication potential of the RNA.

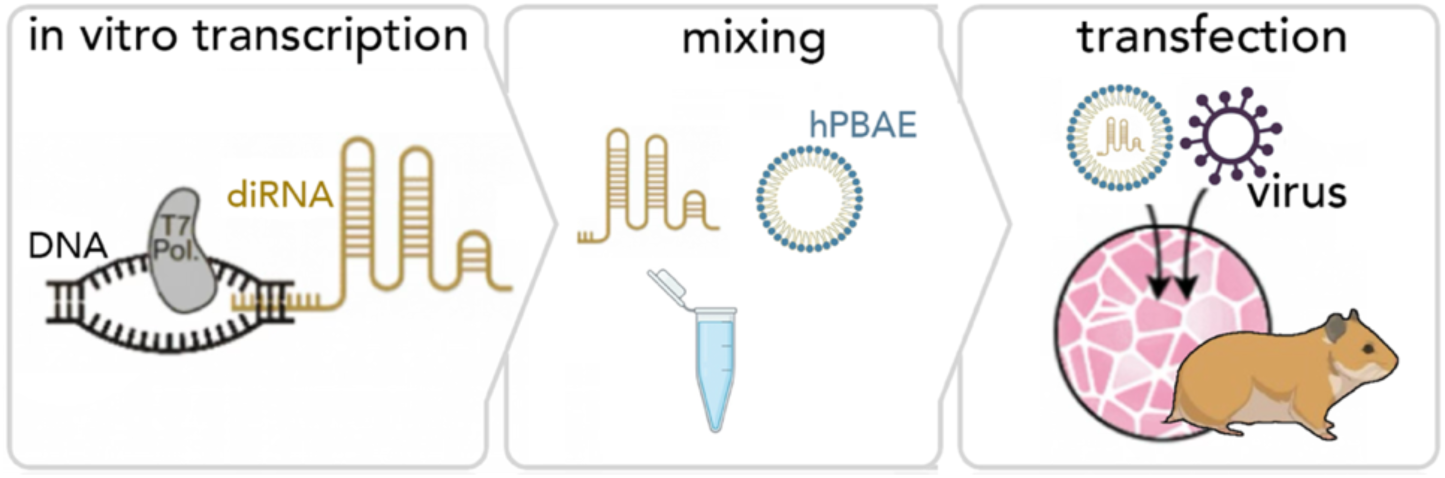

## INTRODUCTION

Lower respiratory tract infections affect approximately 500 million people annually [1], a burden that increased substantially during the coronavirus disease 2019 (COVID-19) pandemic [2]. Vaccines against severe acute respiratory syndrome coronavirus 2 (SARS-CoV-2) were rapidly developed, most notably mRNA vaccines delivered intramuscularly via lipid nanoparticles (LNPs) [3]. In contrast, antiviral development has been slower and less effective. Monoclonal antibodies rapidly lost efficacy against emerging SARS-CoV-2 variants, polymerase and protease inhibitors had limited or transient benefit as resistance evolved, and host-targeted therapies have similarly been constrained by toxicity and modest efficacy [4–6]. More broadly, antiviral discovery remains dependent on drug repurposing or large-scale small-molecule screening, both of which are costly and time-intensive [7,8]. There is therefore a need for antivirals that match the speed, scalability, and adaptability of mRNA vaccines.

Defective interfering RNA (diRNA) represents a potential solution. Delivered to infected cells, diRNA exploits the target virus replication machinery for amplification while suppressing virus replication and transmission [9]. Packaged into virions, diRNA can propagate alongside the virus, promoting attenuation or extinction. This approach has been explored for SARS-CoV-2 [10,11] and offers the potential for rapid, sequence-based antiviral design. However, effective delivery remains a key challenge. Intramuscular LNP delivery is poorly suited for respiratory infections, as it does not efficiently target the lungs. Moreover, LNPs are difficult to aerosolize and may limit therapeutic efficacy to early stages of infection in the upper respiratory tract [11]. Direct pulmonary delivery is therefore desirable, particularly at later stages when viral load peaks in the lungs.

Designing nanoparticles for lung delivery presents multiple constraints. Particles must be small (<300 nm) for mucus penetration and cellular uptake [12–15], yet deliverable as 1–5 μm droplets for optimal lung deposition [16]. Surface charge must balance stability and low inflammation with effective mucus penetration, which favors near-neutral charge [17,18]. Additionally, nanoparticles must withstand nebulization, be non-toxic, be manufacturable at scale, and ideally remain stable without ultra-low-temperature storage [19–21].

Hyperbranched poly(beta-amino ester) (hPBAE) nanoparticles offer a promising platform that addresses many of these challenges. hPBAE form nanoscale polyplexes (a type of nanoparticles) with RNA that facilitate cellular uptake and endosomal escape via pH-responsive release [22–26] and can deliver mRNA [26] and miRNA [27] to lung cells in vivo. However, the delivery of diRNA, which for viruses with a large genome can be several kilobases long, has not been tested, and optimal formulation parameters, including polymer-to-RNA ratios, are unclear. Furthermore, limited data exist on formulation stability, storage conditions, and toxicity of hPBAE-RNA complexes.

We evaluated the potential of hPBAE nanoparticles to deliver diRNA antivirals against betacoronaviruses using multiple formulations and storage conditions. We tested delivery and cellular uptake across diverse cell types, protection against nuclease degradation, and cytotoxicity in vitro and systemic physiological effects in vivo.

## MATERIALS AND METHODS

### Synthesis of hyperbranched poly(beta-amino ester) (hPBAE)

D90−118 hyperbranched poly(beta-amino ester) (hPBAE) was synthesized as previously described (**Supplementary Fig. S1 and S2**) [26,27]. Briefly, acrylate (DD), backbone amine (90), and trifunctional amine monomers were reacted at a molar ratio of 1:0.5:0.2 in anhydrous dimethylformamide (150 mg/mL) at 40 °C for 4 h and then at 90 °C for 48 h. After cooling to 30 °C, end-cap amine (1.5 molar equivalents) was added and the reaction proceeded for 24 h. Polymers were purified by precipitation in cold diethyl ether containing acetic acid, followed by centrifugation at 1,250 × g for 2 min, washing twice in fresh diethyl ether, and vacuum drying for 48 h. Polymer identity was confirmed by nuclear magnetic resonance (NMR) spectroscopy, and samples were stored at −20 °C.

### RNA preparation and in vitro transcription

Cy5-labeled mRNA encoding enhanced green fluorescent protein (*EGFP*) (996 nt) containing 5-methoxyuridine (5-moUTP) in place of uridine and capped with an anti-reverse cap analog was obtained from APExBIO (#R1009). Defective interfering RNAs (diRNAs) for bovine coronavirus (2,765 nt) and SARS-CoV-2 (2,880 nt) [10] were synthesized by in vitro transcription using T7 RNA polymerase and the HiScribe High Yield RNA Synthesis Kit (New England Biolabs, #E2040S). RNA was purified following transcription and assessed for integrity and concentration by automated electrophoresis (Agilent TapeStation). All RNA samples were stored at −80 °C.

### Formation and characterization of hPBAE-RNA nanoparticles

Dimethyl sulfoxide (DMSO) was added to 1 mg of hPBAE and sonicated until dissolved (10-60 sec.); the same volume of sodium acetate (0.1M, pH 5.2) was then added and the solution sonicated again a few seconds to one min, yielding an hPBAE solution at 10 or 50 μg/μL. Unless otherwise specified, RNA was added to hPBAE at a 40:1 (w/w) hPBAE:RNA weight ratio, and mixed by pipetting 10 times. Cryo-transmission electron microscopy (cryo-TEM) was used to confirm that hPBAE–RNA nanoparticles were spherical in morphology. Particle size and ζ-potential were measured in phosphate-buffered saline (PBS) using dynamic light scattering (DLS; Malvern Zetasizer Nano ZS) to verify successful nanoparticles formation and good colloidal stability [21, 26, 27].

### hPBAE–RNA nanoparticle storage and processing

For storage studies, hPBAE and hPBAE–RNA nanoparticles were maintained under various temperature conditions, from room temperature to −80 °C. Lyophilized samples were frozen before drying, and lyophilization was performed using a freeze dryer (Labconco FreeZone 18 Liter Console Freeze Dryer) under standard cycle conditions, followed by reconstitution prior to use. For selected experiments, sucrose (0–90 μg/μL) was added as a cryoprotectant. Aerosolized nanoparticles were produced using an Aeroneb vibrating mesh nebulizer (Aerogen, Kent Scientific, # NEB-7000) to generate droplets in the respirable size range (2.5–4.0 μm according to the manufacturer specifications). Nebulized aerosols were collected and reconstituted in liquid form prior to in vitro application. hPBAE–RNA nanoparticles were added directly to the culture medium at defined doses.

### Cationic lipid–mediated RNA transfection

As an alternative delivery method to hPBAE nanoparticles, synthetic RNA was delivered to cultured cells using MessengerMax (Thermo Fisher Scientific), a cationic lipid–based transfection reagent. RNA–lipid complexes were prepared according to the manufacturer’s instructions by mixing RNA with MessengerMax in Opti-MEM and incubating for 10 min at room temperature prior to transfection. The complexes were then added to cells and incubated under standard culture conditions prior to downstream analyses.

### Cell culture

All cell lines were maintained at 37 °C in a humidified incubator with 5% CO₂ and routinely tested to confirm the absence of mycoplasma contamination. Cells were passaged using enzymatic dissociation at 70–90% confluency and seeded at densities optimized for each experimental assay. Human lung adenocarcinoma A549 cells (ATCC #CCL-185) were cultured in F-12K Nutrient Mixture supplemented with 10% fetal bovine serum (FBS). Human malignant mesothelioma NCI-H28 cells (ATCC #CRL-5820) and human gastric carcinoma NUGC-3 cells (JCRB #0822) were maintained in RPMI-1640 with 10% FBS. High-glucose Dulbecco’s Modified Eagle Medium (DMEM) supplemented with 10% FBS was used for human embryonic kidney HEK293 cells (ATCC #CRL-1573), human hepatocellular carcinoma JHH-7 cells (JCRB #0403), human neuroblastoma SK-N-AS cells (ATCC #CRL-2137), and Vero cells derived from *Chlorocebus sabaeus* kidney epithelium (ATCC #CCL-81). McCoy’s 5A medium supplemented with 15% FBS was used to culture human osteosarcoma SaOS-2 cells (ATCC #HTB-85), whereas McCoy’s 5A medium supplemented with 10% FBS was used to culture human colorectal adenocarcinoma HT-29 cells (ATCC #HTB-38). Eagle’s Minimum Essential Medium (EMEM) supplemented with 10% FBS was used for human hepatocellular carcinoma HepG2 cells (ATCC #HB-8065), human glioblastoma U-87 MG cells (ATCC #HTB-14), human airway epithelial Calu-3 cells (ATCC #HTB-55), human breast adenocarcinoma MCF-7 cells (ATCC #HTB-22), and bovine kidney epithelial MDBK cells (ATCC #CCL-22).

### Cytotoxicity Assay

Cell viability was evaluated using the 3-(4,5-dimethylthiazol-2-yl)-2,5-diphenyltetrazolium bromide (MTT) assay [28]. Cells were seeded in 96-well plates and cultured in DMEM at 37 °C in a humidified atmosphere containing 5% CO₂ for 24 h to reach approximately 70% confluence. Cells were then treated with varying concentrations of nanoparticles or nanoparticle–RNA complexes for 24 h. Following treatment, 10 µL of MTT solution (5 mg/mL in PBS) was added to each well and incubated for 4 h at 37 °C. The medium was carefully removed, and 100 µL of DMSO was added. Plates were shaken for 10 min, and absorbance was measured at 570 nm using a microplate reader.

### Detection of RNA by fluorescence imaging

*EGFP*-positive cells were quantified using a Countess III FL automated cell counter (Thermo Fisher Scientific) following cell detachment, and *EGFP*-positive events were used to estimate transfection efficiency. Cells transfected with Cy5-labeled *EGFP* mRNA were also imaged using a Zeiss Axio Observer fluorescence microscope. Signals from Cy5 (RNA), *EGFP* (protein expression), and 4′,6-diamidino-2-phenylindole (DAPI; nuclei) were acquired using appropriate filter sets. Imaging parameters were kept constant across all samples.

### Detection of RNA by fluorescence in situ hybridization

RNA localization was analyzed using single-molecule fluorescence in situ hybridization (FISH) [29] using Stellaris probes (Custom Assay with CAL Fluor Red 610 Dye, #SMF-1082-5, LGC Biosearch Technologies) [30]. Probe sets targeting diRNA were designed using the Stellaris Probe Designer (www.biosearchtech.com/stellarisdesigner, LGC, Biosearch Technologies, Petaluma, CA) and consisted of 48 singly labeled 20-mer oligonucleotides complementary to non-overlapping regions of the target RNA [31] (masking level: 2; oligo length: 20 nt; minimum spacing length: 2 nt). Fixed and permeabilized cells were hybridized with probe mixtures in hybridization buffer (Stellaris #SMF-HB1-10) containing 10% (v/v) formamide. Following hybridization, samples were washed using Stellaris wash buffers (1:7:2 v/v formamide:water:Buffer A; #SMF-WA1-60; #SMF-WB1-20) to remove unbound probes and reduce background fluorescence. Nuclei were counterstained with 4′,6-diamidino-2-phenylindole (DAPI) at 5 ng/mL in Wash Buffer A. Images were acquired using a widefield fluorescence microscope (Zeiss Axio Observer) with exposure settings kept constant across samples within an experiment.

### Quantification of RNA by RT-qPCR

Total RNA from viral supernatants and infected cells was extracted using TRIzol reagent (Invitrogen) according to the manufacturer’s instructions. Quantification of viral RNA and defective interfering RNA (diRNA) was performed using one-step RT-qPCR with sequence-specific dual-labeled hydrolysis (TaqMan) probes and corresponding primers on a StepOnePlus Real-Time PCR System (Applied Biosystems).

Reactions were performed in triplicate and included no-template controls. RNA copy number in supernatants was quantified using standard curves generated from serial dilutions of in vitro–transcribed RNA. The one-step RT-qPCR reactions were performed using 20 µL reactions containing 1 µg of RNA per sample, 10 µL of FastVirus (Applied Biosystems # 4444434) Master Mix, 0.2 µM of TaqMan probe, and 0.5 µM of primers, under the following cycling conditions: 50 °C for 10 min, 95 °C for 2 min, followed by 40 cycles of 95 °C for 15 s and 60 °C for 1 min.

For RNA in cell extracts, results were normalized using an Eukaryotic 18S rRNA Endogenous Control (FAM/MGB probe, non-primer limited) (Applied Biosystems #4333760F); for a 20 μL reaction, 10 μL TaqMan Fast Universal PCR Master Mix (Applied Biosystems # 4364103), 1 μL primer-probe mix and 100 ng cDNA template. Cycling conditions were 95 °C for 20 s followed by 40 cycles of 95 °C for 3 s and 55 °C for 30 s.

### Propagation of bovine coronavirus and infection in vitro

Bovine Coronavirus Mebus strain was propagated in MDBK cell lines cultured at 37°C and 5% CO_2_ in Eagle’s Minimum Essential Medium (MEM) (CellGro) with 1% GlutaMax I (Gibco) and 10% fetal bovine serum (FBS) (Atlanta Biologicals). The virus was collected and stored in aliquots at −80 °C until the experiments. MDBK cells (80–90% confluent) in 12-well plates were infected at MOI of 1 and incubated at 37 °C in 5% CO₂ for 2 h before the medium was removed, the cells were washed twice with DPBS, and fresh maintenance medium was added.

### Animal studies

Five-week-old golden hamsters (*Mesocricetus auratus*, Waterhouse, 1839) were assigned to three cohorts randomly (36 animals/cohort). In each cohort, 6 hamsters were assigned per treatment group. Hamsters in each group were exposed to either PBS or hPBAE–diRNA intranasally at day zero, and body weight was measured from 10 days before to 5 days after exposure. On day 5, animals were anesthetized with 4% isoflurane in an induction box and blood was collected from each animal at the time of euthanasia. Postmortem examinations were performed and gross findings were documented.

Tissues were collected in BB tubes or trimmed to less than 1-cm thick blocks, placed in cassettes, and fixed for >72 h in 10% neutral-buffered formalin for histopathology. About 5-10% of each lung lobe was collected for histopathology. Microscopic examination of hematoxylin and eosin-stained paraffin sections was performed on tissues collected at necropsy. Observations were scored by severity in compliance with standard guidelines. The Piccolo General Chemistry 13 was used for the quantitative determination of blood markers.

Total RNA was extracted from hamster lungs and other organs using the KingFisher Flex automated extraction system (Thermo Fisher Scientific) in combination with the MagMAX RNA Isolation Kit. Tissues were first homogenized in lysis buffer to disrupt cells and release nucleic acids. Homogenates were then processed on the KingFisher Flex, which uses magnetic bead–based capture to bind RNA, followed by sequential wash steps to remove contaminants. Purified RNA was eluted in RNase-free water and stored at −80°C until downstream applications.

All animal experiments were conducted at the Integrated Research Facility at Fort Detrick (IRF-Frederick), National Institute for Allergy and Infectious Diseases (NIAID), Division of Clinical Research (DCR), National Institutes of Health (NIH) in accordance with the guidelines of the NIH and the Guide for the Care and Use of Laboratory Animals. The IRF-Frederick is accredited by the Association for Assessment and Accreditation of Laboratory Animal Care (AAALAC; 000777), approved for Laboratory Animal Welfare by the Public Health Service (PHS; D16-00602), and registered with the United States Department of Agriculture (USDA; 51-F-0016). The study protocol (IRF-070E) was reviewed and approved by the Institutional Animal Care and Use Committee (IACUC) of the National Institute of Allergy and Infectious Diseases. All procedures were performed in AAALAC International–accredited facilities, and followed the recommendations provided in The Guide for the Care and Use of Laboratory Animals [32], the American Veterinary Medical Association (AVMA) guidelines for the euthanasia of animals [33], and the ARRIVE (Animal Research: Reporting *In Vivo* Experiments) guidelines 2.0 [34]. Efforts were made to minimize animal pain and distress, including the use of isoflurane anesthesia for all terminal procedures.

## RESULTS

### Efficient intracellular delivery of RNA in lung epithelial cells

To study the efficiency of hPBAE nanoparticles as delivery vectors, we started by verifying intracellular delivery of Cy5-labeled *EGFP* mRNA to A549 cells. Confocal microscopy revealed both Cy5 signal and *EGFP* expression 12 h after transfection with hPBAE nanoparticles carrying *EGFP* mRNA, confirming intracellular delivery of functional mRNA (**Fig. 1A**).

**Figure 1.**
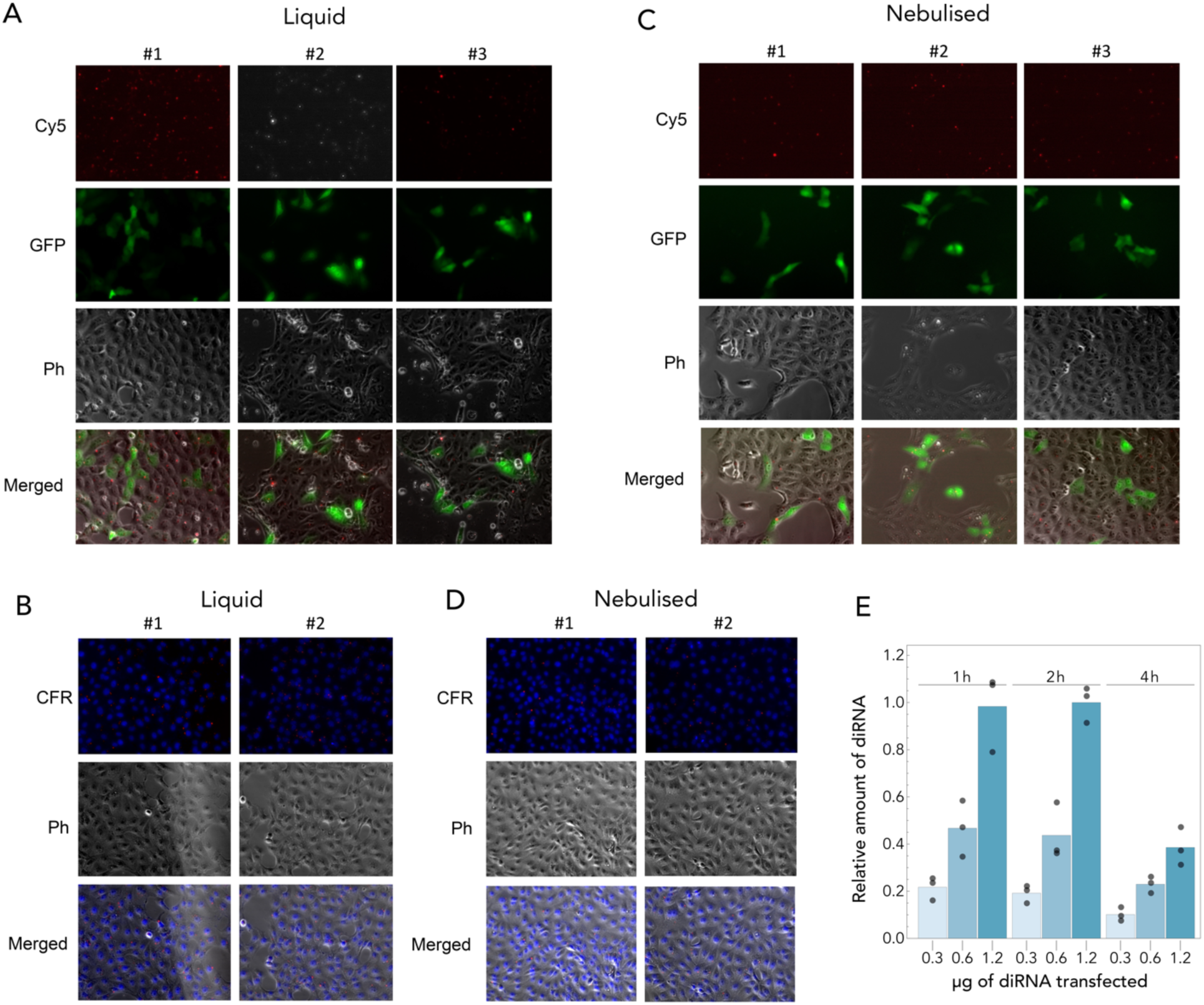
Efficient intracellular delivery of hPBAE–RNA nanoparticles into lung epithelial cells. Cy5-labeled *EGFP* mRNA (**A, C**) or diRNA targeting SARS-CoV-2 (**B, D, E**) were delivered to A549 cells using hPBAE nanoparticles either in solution (**a, b**) or following nebulization and reconstitution in liquid form (**C, D**). Cells were imaged by confocal microscopy 12 h post-transfection. For hPBAE–mRNA nanoparticles (**A, C**) Cy5 (red) indicates RNA localization, and *EGFP* (green) indicates expression. diRNA (**B, D**), was detected by FISH, with CAL Fluor Red 610 (CFR, red) indicating RNA localization. In all panels, Ph denotes phase-contrast imaging to show cell morphology, and nuclei were stained with 4′,6-diamidino-2-phenylindole (DAPI, blue). **E:** hPBAE–diRNA nanoparticles (delivered without nebulization) were quantified by RT-qPCR in cell lysates extracted after 1, 2 or 4 h post-transfection.

Next, we replaced the 996-nt long *EGFP* mRNA with 2,880 nt long SARS-CoV-2 diRNA. hPBAE nanoparticles carrying this diRNA had a diameter of 75±16 nm (**Supplementary Fig. S3**), and a *ζ*-potential of 23.89±0.56 mV, indicating successful nanoparticle formation and diRNA loading.

Delivery was confirmed by fluorescence in situ hybridization (FISH), which showed robust intracellular signal 12 h after transfection (**Fig. 1B**). RT-qPCR detected ∼45 ng of intracellular diRNA 1 h after delivery of a 500-ng dose (a ∼90% reduction, likely reflecting incomplete delivery and losses during extraction), with levels decreasing over time (approximately 50% from 1 h to 4 h post-transfection). No intracellular diRNA was detected in the absence of hPBAE, indicating that uptake was carrier-dependent.

To assess robustness to aerosolization, both types of nanoparticles were nebulized prior to transfection. Nebulized samples could be delivered efficiently, as evidenced by mRNA expression (**Fig. 1C**) and diRNA detection (**Fig. 1D**)

Delivery efficiency was dose-dependent, with intracellular diRNA levels proportional to the amount loaded onto nanoparticles (**Fig. 1E**). Together, these results demonstrate efficient cytosolic delivery of functional 996 nt mRNA or 2880 nt-long diRNA by hPBAE nanoparticles into lung epithelial cells, and that hPBAE-based nanoparticles remain functional after shear stress caused by nebulization.

### Optimization of the hPBAE:RNA formulation

To improve payload delivery, we optimized the hPBAE:RNA ratio. Delivery efficiency increased with hPBAE content, with hPBAE:RNA ratios ≥40:1 (w/w) yielding optimal performance for both hPBAE–mRNA and hPBAE–diRNA formulations (**Fig. 2A,B**).

**Figure 2.**
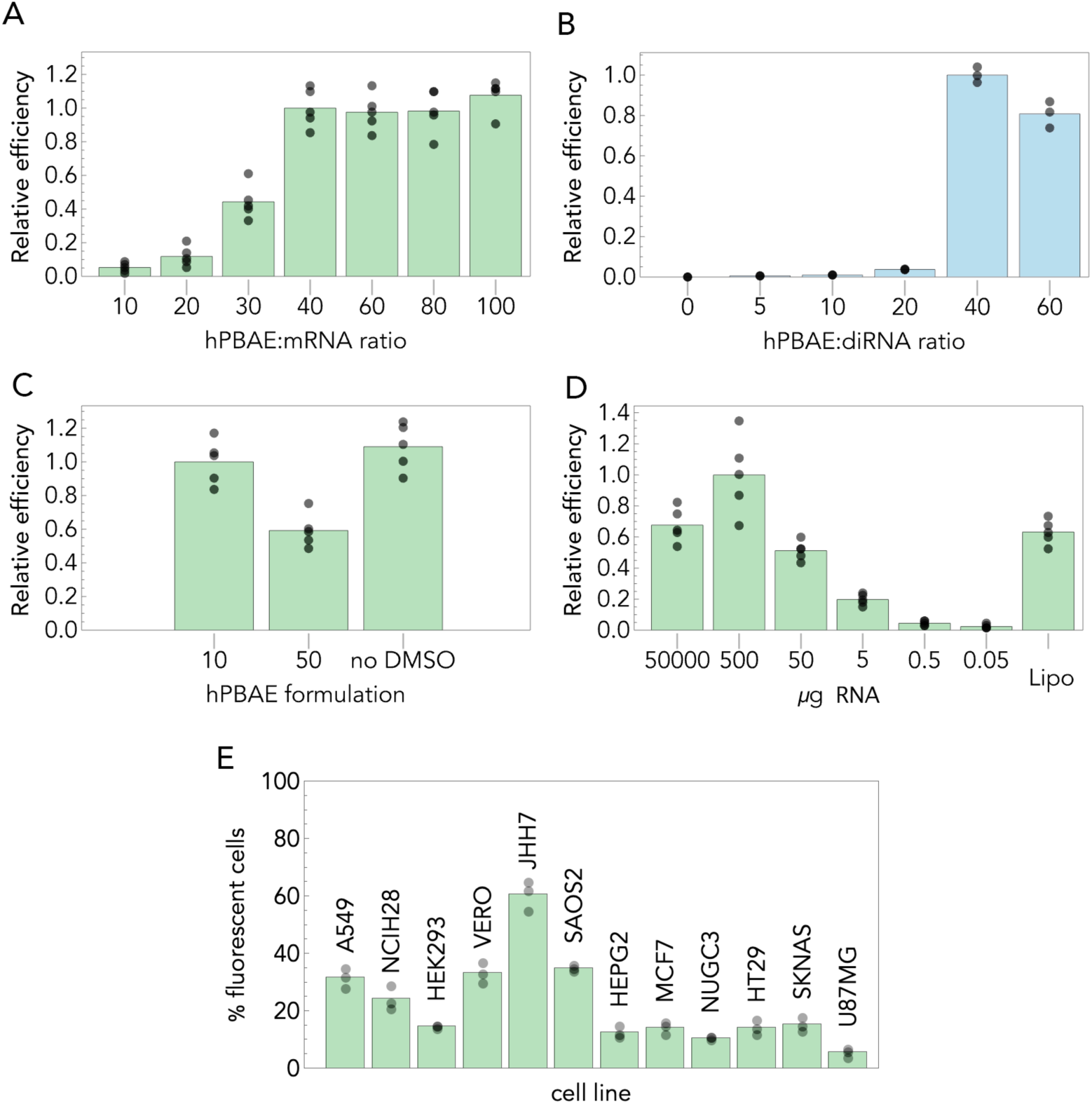
Optimization of the hPBAE–RNA formulation. *EGFP* mRNA (**A**) or SARS-CoV-2 diRNA (**B**) were delivered to Vero cells using hPBAE nanoparticles at varying hPBAE:RNA ratios. Transfection efficiency was quantified as the fraction of fluorescent cells (for *EGFP* mRNA) or by RT-qPCR (for diRNA). **C:** *EGFP* mRNA was delivered to Vero cells using hPBAE prepared under different solvent conditions: dimethyl sulfoxide (DMSO) and sodium acetate in equal volumes; at a concentration of hPBAE of 10 or 50 μg/μL; or sodium acetate only (no DMSO) at a concentration of hPBAE of 20 μg/μL. Transfection efficiency was estimated by the fraction of fluorescent cells. **D**: Increasing amounts of *EGFP* mRNA were delivered to Vero cells using hPBAE nanoparticles, and transfection efficiency was quantified as the fraction of fluorescent cells. A cationic lipid–based transfection reagent (Lipo) was included for comparison. **E:** *EGFP* mRNA was delivered to different cell lines using hPBAE nanoparticles, and transfection efficiency was quantified by the fraction of fluorescent cells. Unless otherwise stated, hPBAE was used at 10 μg/μL in equal volumes of DMSO and sodium acetate, and 500 ng of RNA was delivered at a 40:1 (w/w) hPBAE:RNA ratio.

Because hPBAE is very viscous, it can be dissolved in DMSO before dilution in an equal volume of sodium acetate, followed by mixing with RNA. We found that a concentration of 10 μg/μL outperformed 50 μg/μL and that pre-dissolution in DMSO prior to dilution in sodium acetate was not essential (**Fig. 2C**).

We next used hPBAE prepared in equal volumes of DMSO and sodium acetate at 10 μg/μL to transfect varying amounts of RNA at a 40:1 hPBAE:RNA weight ratio into the same number of cells. Delivery of 500 ng of *EGFP* mRNA to approximately 4 × 10⁵ cells (equivalent to a confluent well of a 12-well plate) resulted in transfection of ∼50% of cells, as assessed by fluorescence microscopy 24 h post-transfection (**Fig. 2D**). Cell density had minimal impact on delivery efficiency, with similar transfection rates observed at 60% confluence compared to full confluence (data not shown). Transfection efficiency varied substantially between hPBAE batches, ranging from 30% to 90%. Reducing the RNA dose (while keeping the same hPBAE:RNA ratio of 40:1) decreased the fraction of transfected cells, while higher doses did not improve delivery and could be counterproductive (**Fig. 2D**).

Finally, delivery efficiency varied across cell types, with lower transfection observed in U-87 MG cells and higher efficiency in JHH-7 cells (**Fig. 2E**), indicating cell-type-dependent uptake.

### Storage, stability, and transfection efficiency of hPBAE–RNA nanoparticles

We evaluated the stability of hPBAE–RNA nanoparticles under different storage conditions. First, hPBAE was stored at 4 °C for up to 24 months, then mixed with *EGFP* mRNA, and the resulting nanoparticles were used for transfection. Storage at 4 °C for up to 24 months resulted in a modest reduction in transfection efficiency (**Fig. 3A**). Although long-term storage at room temperature was not systematically assessed, storage for up to one week did not impair transfection efficiency.

**Figure 3.**
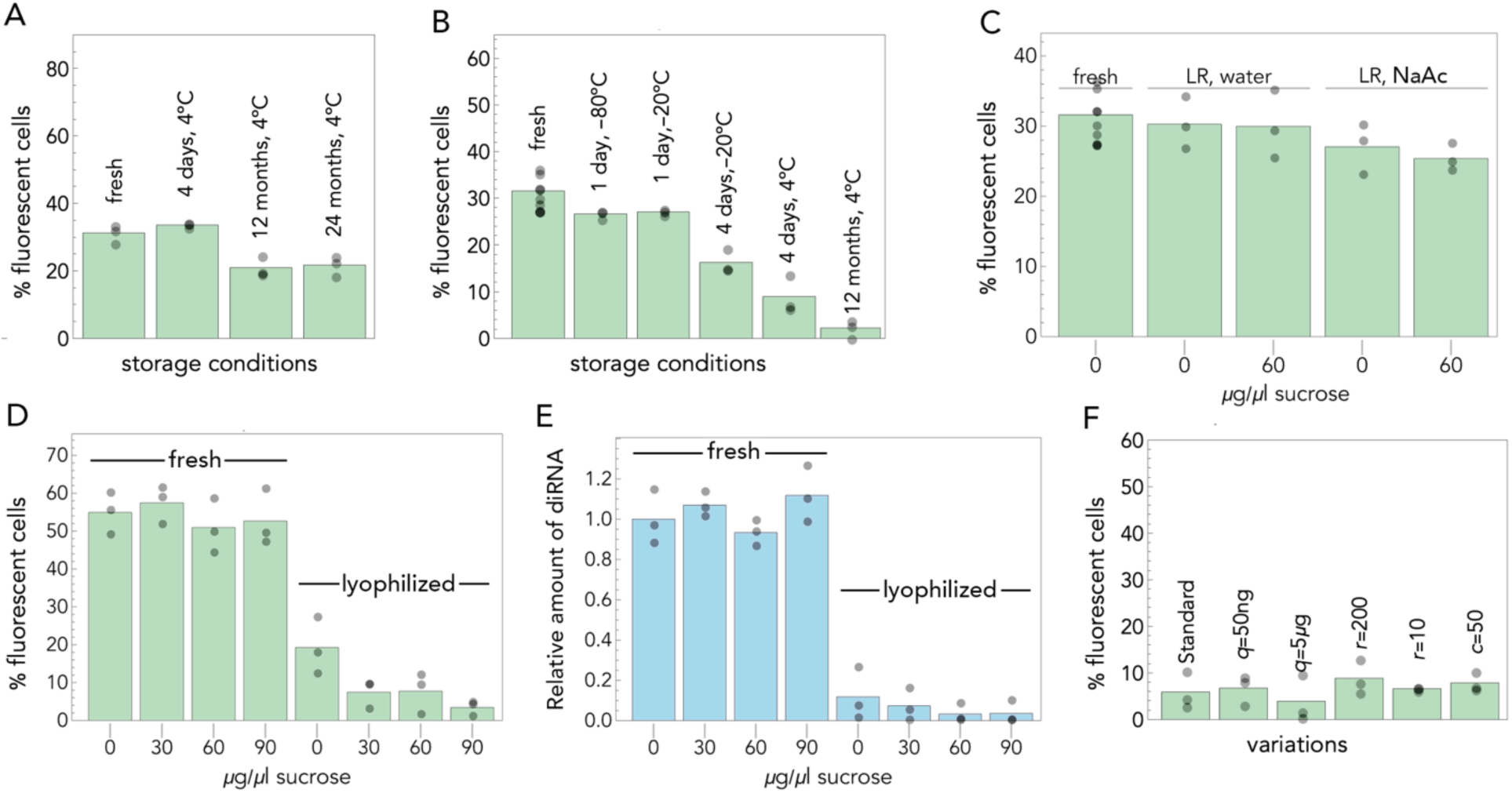
Storage, stability, and transfection efficiency of hPBAE–RNA nanoparticles. hPBAE was mixed with *EGFP* mRNA (green) or diRNA targeting SARS-CoV-2 (blue), and transfection efficiency was assessed in Vero cells as the fraction of fluorescent cells at 24 h post-transfection (*EGFP* mRNA) or by RT-qPCR at 4 h post-transfection (diRNA). **A**: hPBAE was stored at 4 °C for varying durations, then mixed with *EGFP* mRNA, and the resulting nanoparticles were used for transfection. **B**: hPBAE was mixed with *EGFP* mRNA, and the resulting nanoparticles were stored under different conditions prior to transfection. **C**: hPBAE was mixed with *EGFP* mRNA with or without sucrose, then lyophilized and reconstituted (“LR”) in water or sodium acetate (“NaAc”). **D**: Fresh or lyophilized hPBAE–mRNA nanoparticles containing varying amounts of sucrose were frozen at −80 °C, then reconstituted in water. **E**: Fresh or lyophilized hPBAE–diRNA nanoparticles containing varying amounts of sucrose were frozen at −80 °C, then reconstituted in water. **F**: Lyophilized hPBAE–mRNA nanoparticles with variations in formulation parameters (q, RNA amount; r, hPBAE:RNA ratio; c, hPBAE concentration in μg/μL) were frozen at −80 °C, then reconstituted in water. Unless otherwise stated (panel f), hPBAE was used at 10 μg/μL in equal volumes of DMSO and sodium acetate, and 500 ng of RNA was delivered at a 40:1 (w/w) hPBAE:RNA ratio.

Next, hPBAE was mixed with *EGFP* mRNA, and the resulting nanoparticles were stored under different conditions prior to transfection. In contrast to hPBAE alone, hPBAE–RNA nanoparticles were less stable, with significant reductions in transfection efficiency after storage at 4 °C or freezing for several days (**Fig. 3B**). Freezing for one day did not significantly affect transfection efficiency; however, freezing or storage at 4 °C for up to four days reduced efficiency by more than 50%. Nanoparticles-RNA stored for 12 months at 4 °C retained limited activity.

Lyophilization followed by immediate reconstitution preserved delivery efficiency, whereas lyophilization combined with freezing substantially impaired delivery. The addition of sucrose or other modifications did not restore this loss (**Fig. 3C–F**).

To assess RNA stability, *EGFP* mRNA was incubated with RNase for 30 min, purified, and then complexed with hPBAE prior to transfection. This resulted in a marked reduction in *EGFP*-positive cells (<1%), indicating near-complete RNA degradation. In contrast, addition of RNase after nanoparticle formation did not significantly reduce *EGFP* expression, indicating that RNA within hPBAE nanoparticles is protected from RNase degradation.

Together, these results indicate that hPBAE can be stored at 4 °C for up to two years after dilution in DMSO and sodium acetate with limited loss of efficacy, and that hPBAE nanoparticles protect RNA from RNase degradation. However, hPBAE–RNA nanoparticles should be used as soon as possible after preparation, preferably on the same day.

### hPBAE is not toxic and delivers functional diRNA in vitro and in vivo

To assess toxicity, bovine coronavirus (BCoV) diRNA was delivered to A549, Vero, and Calu-3 cells using hPBAE at 10 μg/μL and a 40:1 (w/w) hPBAE:RNA ratio. No significant cytotoxicity was observed except at very high RNA doses (600 ng) (**Fig. 4A**). Next, cells were infected with BCoV at a MOI of 1 followed, 1 h post-infection, by transfection with hPBAE nanoparticles carrying BCoV diRNA. Cells and supernatants were collected at 1 h and 24 h post-infection to quantify BCoV RNA and diRNA levels. Robust diRNA replication was observed in BCoV-infected cells but not in uninfected controls, confirming that the delivered diRNA is replication-competent in the presence of the helper virus (**Fig. 4B**). diRNA levels increased in both cell lysates and supernatants, indicating intracellular amplification and packaging into BCoV particles.

**Figure 4.**
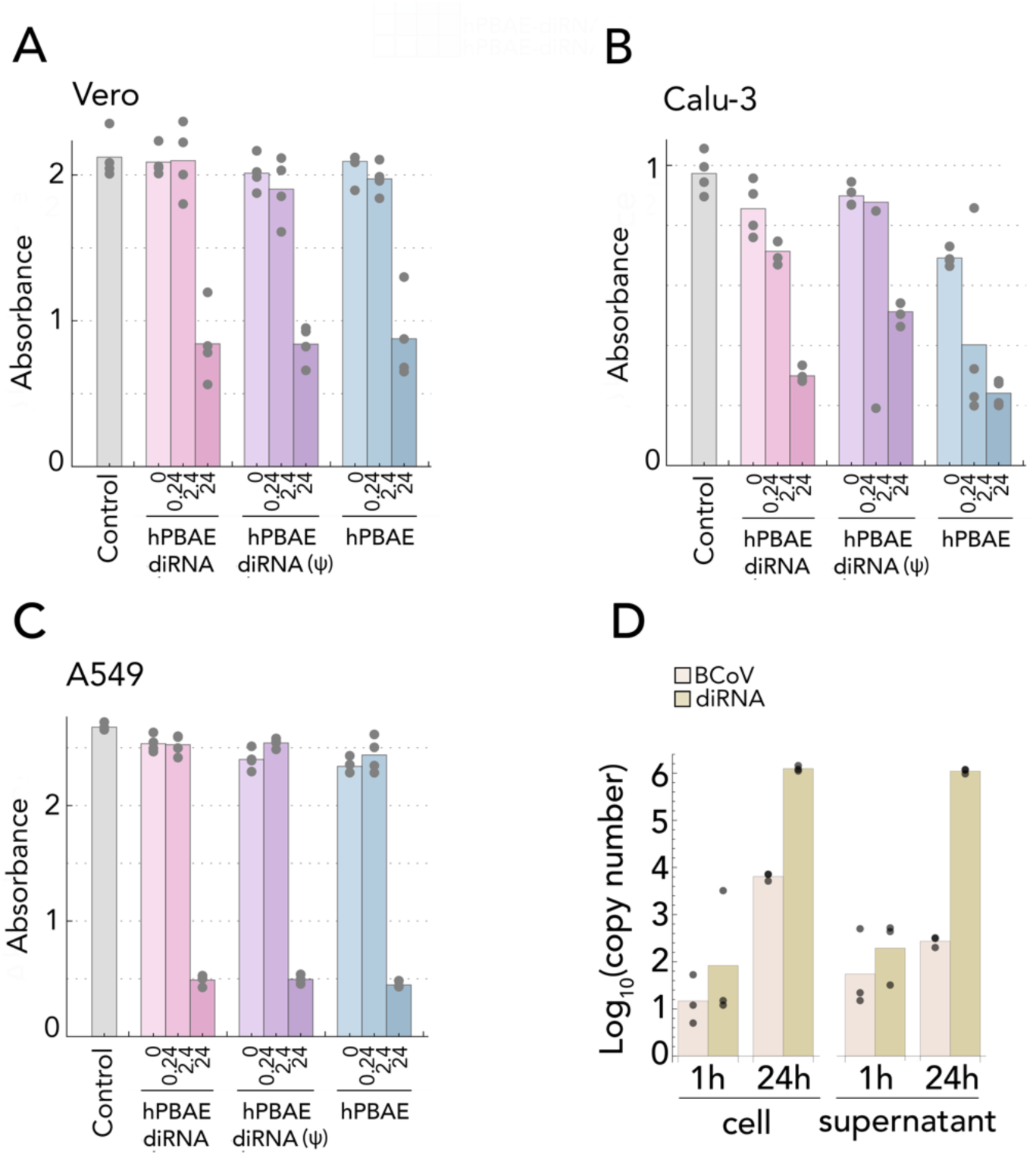
hPBAE nanoparticles are not toxic and deliver functional diRNA in vitro. **A**: 0.24, 2.4 or 24 μg hPBAE, with or without diRNA (at a 40:1 hPBAE:RNA weight ratio), with or without pseudouridin modification (ψ), were delivered to (**A**) Vero, (**B**) Calu-3, or (**C**) A549 cells and cell viability was measured by MTT assay. **D**: hPBAE–diRNA nanoparticles (0.3 μg of RNA) were delivered to MDBK cells 1 h after exposure to BCoV (MOI of 1), and the amount of BCoV genomic RNA and diRNA in cell extracts and in supernatants were measured after 1 or 24 h.

We next evaluated in vivo delivery in golden hamsters. Intranasal administration (three 50 μL doses, 12 h apart) of hPBAE carrying SARS-CoV-2 diRNA resulted in measurable levels of diRNA in the lungs of treated animals after 4 days, whereas diRNA delivered without hPBAE was not detected (**Fig. 5A**). These results indicate that diRNA can be effectively delivered in vivo to the lungs via intranasal administration of hPBAE nanoparticles, even in the absence of nebulization. Treatment was well tolerated in hamsters, with no significant differences in body weight (measured from 10 days pre-treatment to 5 days post-treatment) observed between treated and control animals throughout the study (**Fig. 5B**). Histological analysis revealed no abnormalities in lung tissue (**Supplementary Fig. S4**) and no detectable changes in liver or brain tissues (data not shown). Blood chemistry analysis revealed no significant differences in markers of liver function, renal function, or metabolic status between treated and control groups (**Table 1**). Together, these findings indicate that hPBAE–diRNA nanoparticles enable effective in vivo delivery without detectable impairment of liver or renal function, and without disruption of metabolic homeostasis.

**Figure 5.**
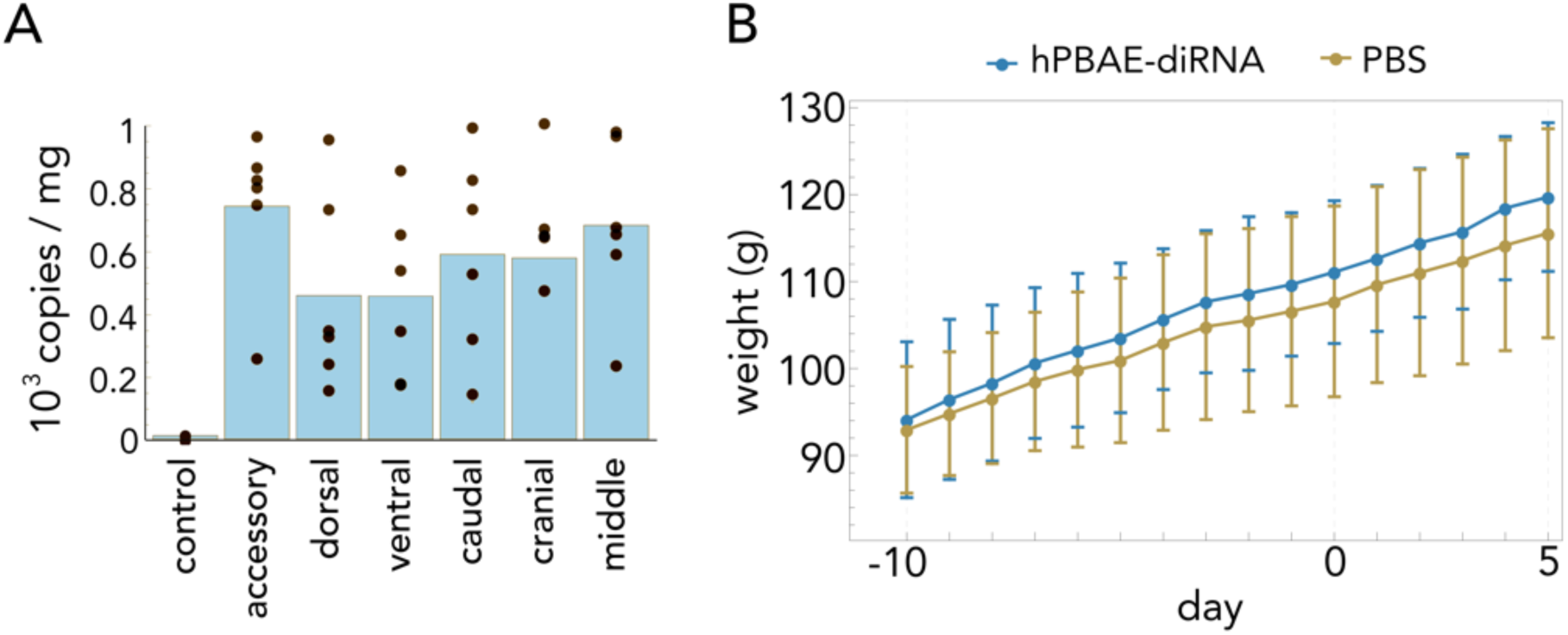
hPBAE nanoparticles are not toxic and deliver diRNA to the lungs. **A**: Golden hamsters were treated with 3 μg of SARS-CoV-2 diRNA and diRNA levels were quantified in different lung regions, 5 days post-treatment. Lung parts are defined as described in standard golden hamster lung anatomy terms [35]. The control treatment was RNA without hPBAE (averaged across lung parts). **B.** The body weights of hamsters, treated with hPBAE–diRNA nanoparticles or mock-treated with phosphate-buffered saline (PBS), measured daily before and after treatment at day 0.

**Table 1.** Changes in the levels of biomarkers in the blood of hamsters treated with hPBE-diRNA polyplexes. *U* statistic and *p* value for the Mann-Whitney U test between treated (hPBAE-diRNA) and non-treated (PBS) hamsters.

| **Marker** | ***U*** | ***p*** |
| --- | --- | --- |
| Alanine aminotransferase | 13 | 0.662 |
| Alkaline phosphatase | 12.5 | 0.537 |
| Aspartate aminotransferase | 13.5 | 0.792 |
| Amylase | 12 | 0.537 |
| Blood urea nitrogen | 10 | 0.548 |
| Glucose | 12 | 0.537 |
| Calcium | 18 | 0.537 |

## DISCUSSION

We have demonstrated that hyperbranched poly(beta-amino ester) (hPBAE) nanoparticles can deliver long, functional diRNA in vitro and in vivo, and remain robust after nebulization. Importantly, we observed no evidence of cytotoxicity in vitro, nor any body weight changes, lung pathology, or alterations in blood chemistry in exposed golden hamsters, supporting the safety of this delivery platform.

It is possible that the detected diRNA originated from viral particles generated in the nasal epithelium following hPBAE–diRNA delivery, rather than from replication of diRNA delivered directly to the lungs via nanoparticles. However, several observations support functional delivery to the lungs: diRNA was detectable five days post-administration, considerably longer than typical mammalian mRNA half-lives (∼7–16 h) [36,37], and previous work has demonstrated efficient pulmonary delivery of hPBAE nanoparticles [26]. Nonetheless, future studies using replication-specific markers or labeled diRNA would further strengthen this conclusion.

Regarding nebulization, our experiments indicate that the nanoparticles resist shear forces; however, we did not assess true aerosol deposition efficiency. Therefore, if an inhalable aerosol formulation is desired, direct delivery experiments without reconstitution will be required.

Cytotoxicity of nanoparticles was observed only at high doses—600 ng of diRNA formulated with hPBAE at a 40:1 ratio, or hPBAE alone, per well in a 96-well plate (∼0.04–0.2 ng per cell, assuming 3,000–15,000 cells per well). These doses are several orders of magnitude above the anticipated therapeutic dose in golden hamsters, indicating negligible toxicity under treatment conditions. For context, the maximum intranasal RNA dose using our standard hPBAE formulation is 10 μg (20 μL). Adult golden hamster lungs (∼1–1.5 g; ∼0.8–1.2% of body mass) are estimated to contain ∼10⁹ cells. Assuming perfectly homogeneous distribution, this would correspond to ∼10⁻⁵ ng RNA per cell—over four orders of magnitude below the cytotoxic levels observed in vitro. Together, these calculations suggest that hPBAE–diRNA nanoparticles are effectively non-cytotoxic at therapeutic doses.

Our in vivo safety assessment reinforces this conclusion. High-throughput blood chemistry analysis revealed no significant deviations in markers of hepatic, pancreatic, renal, endocrine, or mineral balance function. These biomarkers serve as indicators of organ and metabolic function. Specifically, alanine aminotransferase (ALT) and aspartate aminotransferase (AST) are intracellular enzymes released into the bloodstream upon hepatocyte damage; AST may also reflect injury to cardiac or skeletal muscle. Elevated levels are typically associated with hepatocellular injury [38]. Alkaline phosphatase (ALP) activity increases primarily in conditions involving cholestasis or biliary obstruction [39]. Amylase activity is elevated in acute pancreatitis, although increases may also result from salivary gland disorders or impaired renal clearance [40]. Blood urea nitrogen (BUN) reflects renal function and protein metabolism; elevated levels may indicate reduced glomerular filtration, hypovolemia, or increased protein catabolism, whereas decreased levels can occur in severe liver dysfunction [41]. Serum glucose reflects carbohydrate metabolism and endocrine function, with hyperglycemia indicating insulin deficiency, insulin resistance, or stress, and hypoglycemia resulting from excessive insulin activity, adrenal insufficiency, or critical illness [42]. Calcium levels reflect parathyroid and renal regulation; hypercalcemia is often associated with hyperparathyroidism, whereas hypocalcemia may arise from hypoparathyroidism or renal failure [43].

Since none of these biomarkers were elevated or significantly deviated from normal ranges, there is no biochemical evidence of acute hepatic injury (ALT, AST), cholestasis or biliary obstruction (ALP), pancreatic inflammation (amylase), impaired renal excretory function or altered protein catabolism (BUN), disturbances in glucose homeostasis (glucose), or abnormalities in calcium regulation involving parathyroid, renal, or bone physiology (calcium). Although normal values do not exclude early or subclinical disease, this pattern supports preserved hepatic, pancreatic, renal, endocrine, and mineral balance physiology, and suggests that major acute pathological processes affecting these organ systems are unlikely at the time of testing. Nevertheless, several limitations should be considered. First, transient or low-level changes in these biomarkers may have been missed if fluctuations occurred outside the sampling window or remained below assay sensitivity. Second, the selected biomarker panel does not capture all parameters relevant to hepatic, renal, metabolic, or pancreatic function, and therefore may not reflect more subtle physiological alterations. Finally, the short observation period (5 days) limits the ability to detect delayed, cumulative, or longer-term effects that may emerge beyond the monitoring window.

The collective in vitro and in vivo findings provide a consistent safety profile. The absence of cytotoxicity in cultured cells, together with the lack of body-weight loss in hamsters, indicates that the treatment did not induce overt toxicity at either the cellular or organismal level. In addition, the absence of lung pathology and the stability of key blood chemistry markers—reflecting renal and hepatic function as well as overall metabolic homeostasis—support the conclusion that no significant physiological stress or organ-specific toxicity occurred. Taken together, these results indicate that under the conditions tested, the intervention was well tolerated and did not produce measurable adverse effects.

In summary, we demonstrate that hyperbranched PBAE (hPBAE) nanoparticles effectively deliver long, functional defective interfering RNA (diRNA) both in vitro and in vivo. hPBAE–diRNA nanoparticlesmaintained RNA integrity following storage, nebulization, and standard handling procedures, and enabled efficient cytoplasmic delivery across multiple cell types as well as to the lungs of golden hamsters. Both in vitro cytotoxicity assays and comprehensive in vivo analyses—including body weight monitoring, lung histopathology, and blood chemistry—showed no evidence of acute toxicity under the tested conditions. These findings establish hPBAE nanoparticles as safe and effective vectors for lung-targeted delivery of long RNA therapeutics, including replication-competent diRNAs against respiratory RNA viruses. Our work provides a foundation for further optimization of RNA formulations and supports the potential application of hPBAE-based delivery systems in antiviral strategies and other RNA-based therapies. Future studies should focus on aerosolized delivery efficiency, long-term safety, and therapeutic efficacy in disease models, highlighting the translational potential of hPBAE nanoparticles for RNA-based interventions in respiratory infections.

## ACKNOWLEDGEMENTS

We acknowledge the Huck Institutes’ Genomics Core Facility (RRID:SCR_023645) for sequencing and we thank the National Institutes of Health (NIH) National Institute of Allergy and Infectious Diseases (NIAID) Division of Clinical Research (DCR) Integrated Research Facility at Fort Detrick (IRF-Frederick) Comparative Medicine and Clinical Core staff for successful implementation of the animal studies.

This project was supported by the Huck Institutes of the Life Sciences at the Pennsylvania State University through the Huck Innovative and Transformational Seed Grant (HITS) and research agreement SARS-CoV-2-HAM-070E-1 between the Pennsylvania State University and the NIH. This work was supported in part through a Laulima Government Solutions, LLC, prime contract with the U.S. National Institute of Allergy and Infectious Diseases (NIAID) under Contract No. HHSN272201800013C. J.A. and G.W. performed this work as employees of Laulima Government Solutions, LLC. J.H.K. performed this work as an employee of Tunnell Government Services (TGS), a subcontractor of Laulima Government Solutions, LLC, under Contract No. HHSN272201800013C. This research was supported in part by the Intramural Research Program of the National Institutes of Health (NIH). The contributions of the NIH author(s) are considered Works of the United States Government. The findings and conclusions presented in this paper are those of the author(s) and do not necessarily reflect the views of the NIH or the U.S. Department of Health and Human Services.

The funders had no role in the design of the study; in the collection, analyses, or interpretation of data; in the writing of the manuscript; or in the decision to publish the results.

## ETHICAL APPROVAL

All animal experiments were conducted in accordance with the National Institutes of Health guidelines and the Guide for the Care and Use of Laboratory Animals. The study protocol was reviewed and approved by the Institutional Animal Care and Use Committee (IACUC) of the National Institute of Allergy and Infectious Diseases (NIAID). All procedures were performed in AAALAC International–accredited facilities. Efforts were made to minimize animal suffering and reduce the number of animals used.

## DATA AVAILABILITY STATEMENT

The data that support the findings of this study are available from the corresponding author, M.A., upon request.

## AUTHOR CONTRIBUTION STATEMENT

SY: synthetized and validated constructs, performed in vitro experiments, analysed data

JA: performed in vivo experiments and analysed data

PD: performed in vitro experiments

IP: performed in vitro experiments

JHA: performed nanoparticle characterization

AKS: synthetized nanoparticles

KG: synthetized nanoparticles

DH: supervised nanoparticle characterization

GW: designed in vivo experiments

JK: designed in vivo experiments, revised the manuscript

MA: designed the study, performed in vitro experiments, analyzed and interpreted data, wrote the manuscript

## SUPPLEMENTARY MATERIAL

**Figure S1.**
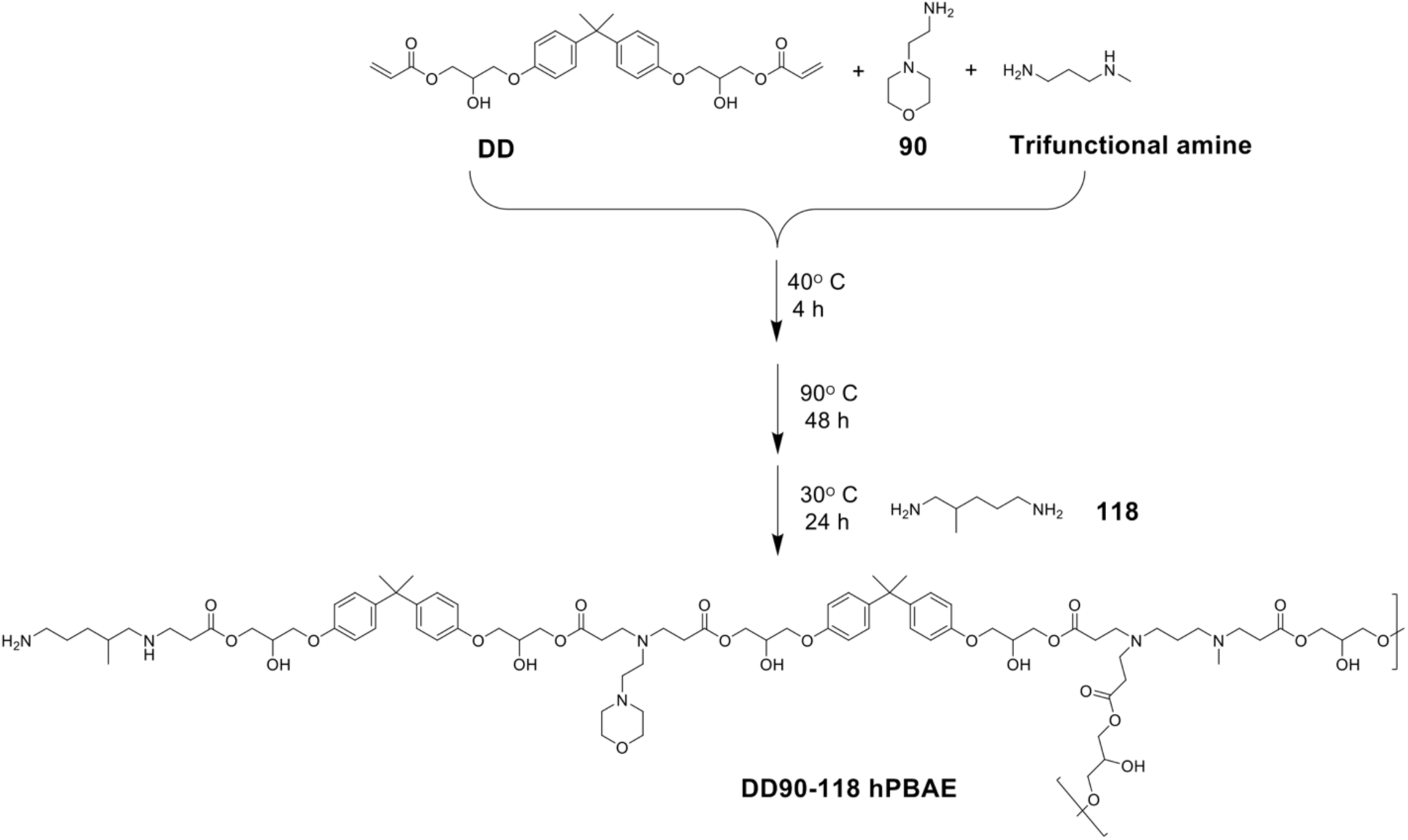
Synthesis of hPBAE. A summary of the process used to produce hPBAE. **b**: Proton Nuclear Magnetic Resonance spectrum of the resulting hPBAE.

**Figure S2.**
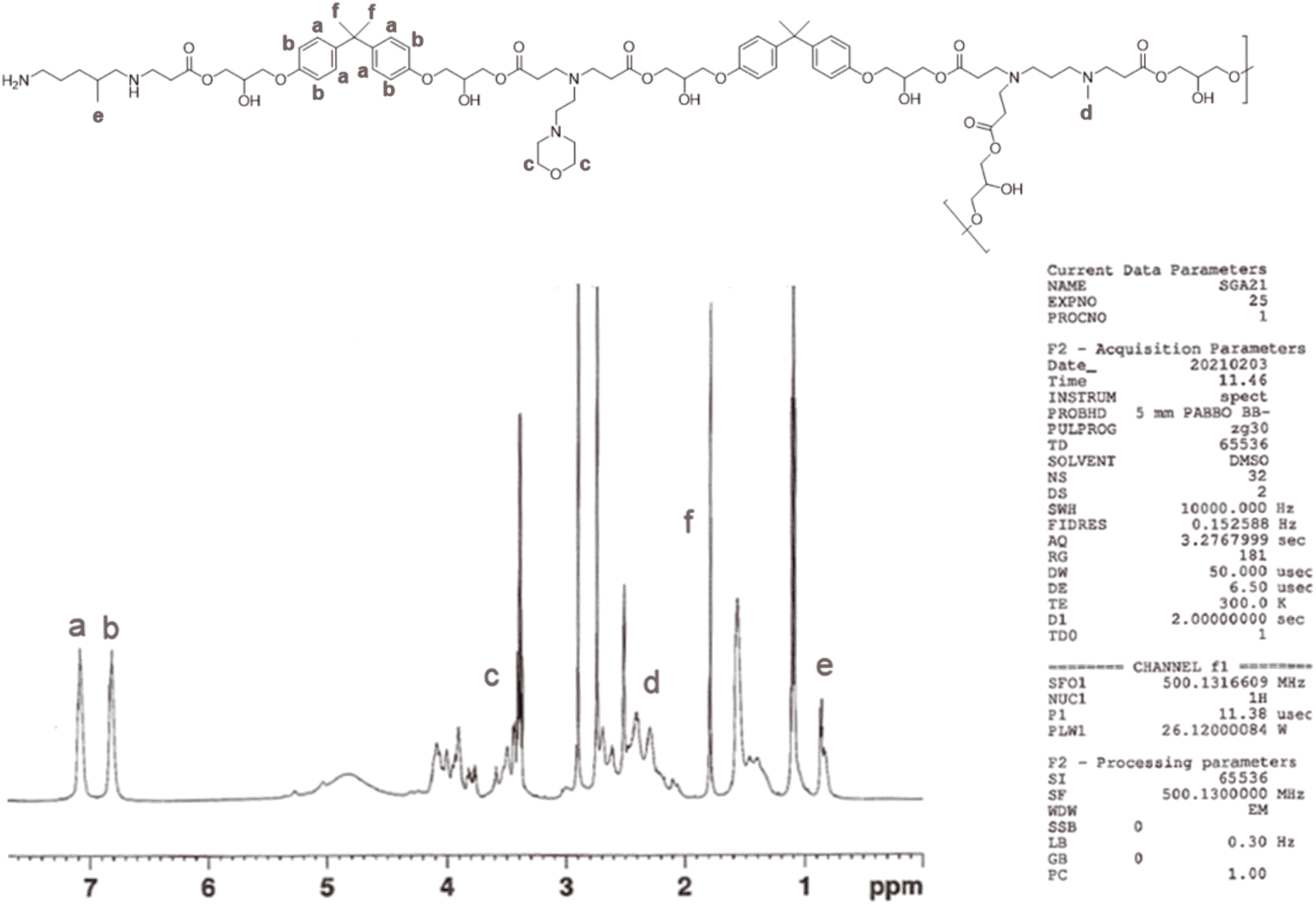
hPBAE synthesis. Proton Nuclear Magnetic Resonance spectrum of the hPBAE used in the study.

**Figure S3.**
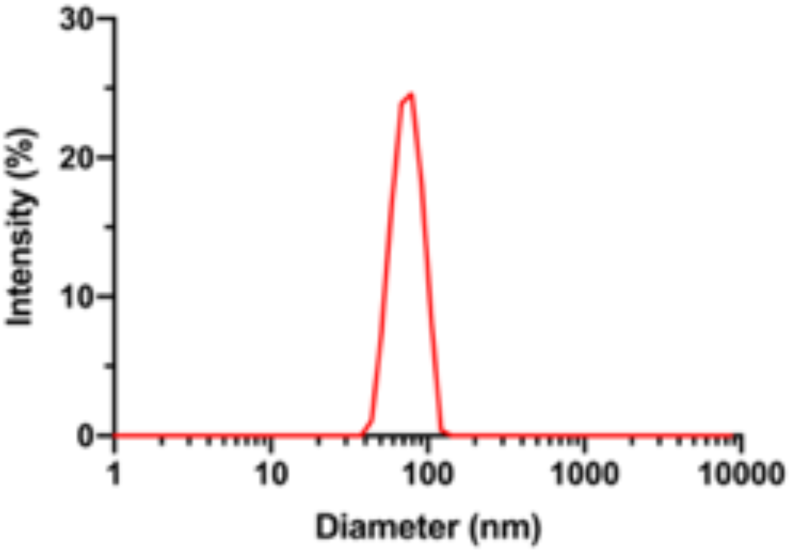
Nanoparticle size. Nanoparticle size distribution of hPBAE-RNA complexes measured by dynamic light scattering.

**Figure S4.**
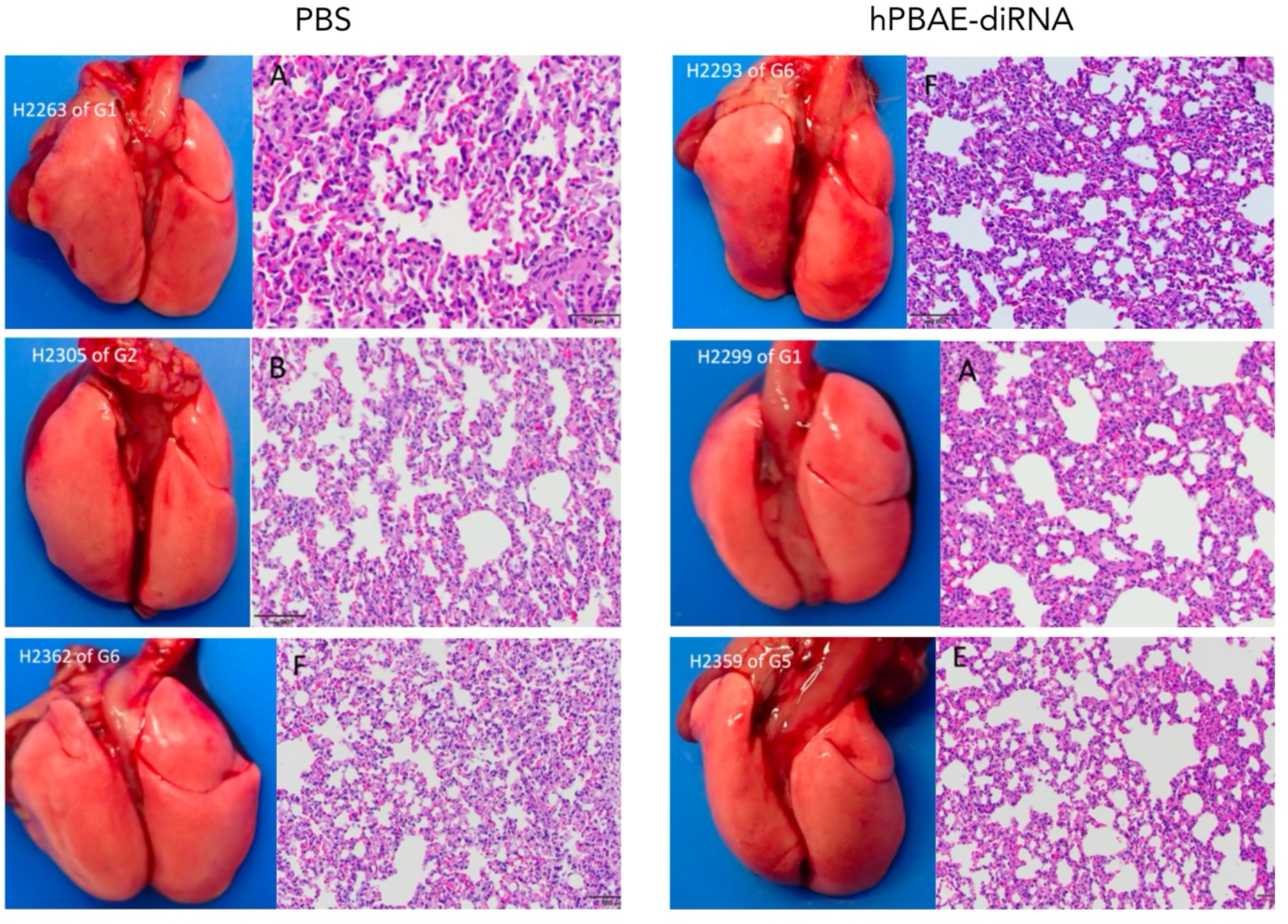
Effect of hPBAE-diRNA on lung pathology. Absence of gross pathology and histopathology in lungs of golden hamsters treated with PBS control or hPBAE-diRNA. The labels shown in the figure correspond to identifiers used during blinded data collection and are not relevant to the interpretation of the results.

